# Species-specific InDel markers for authentication of the Korean herbs *Zanthoxylum schinifolium* and *Zanthoxylum piperitum*

**DOI:** 10.1101/646240

**Authors:** Yonguk Kim, Jawon Shin, Seung-Sik Cho, Yong-Pil Hwang, Chulyung Choi

**Affiliations:** Jeonnam Institute of Natural Resources Research, Jangheung-gun, Jeollanam-do 59339, Korea; Department of Pharmacy, College of Pharmacy, Mokpo National University, Muan-gun, Jeollanam-do 58554, Korea; Department of Pharmaceutical Engineering, International University of Korea, Jinju-si, Gyeongsangnam-do 52833, Korea

**Keywords:** chloroplast genome, chopi, InDel markers, nucleotide diversity, sancho, *Zanthoxylum piperitum*, *Zanthoxylum schinifolium*

## Abstract

**Objective:** *Zanthoxylum schinifolium* and *Zanthoxylum piperitum* are the sources of the well-known traditional Korean herbal medicines “sancho” (prickly ash) and “chopi” (Korean pepper), respectively. Sancho and chopi are often indiscriminately mixed due to the similar appearance of the herbal materials when used as spices and herbal medicines. Moreover, commercial sancho and chopi products often contain adulterants, which could compromise drug efficacy and safety.

**Methods:** In this study, we developed hypervariable InDel markers to distinguish between sancho and chopi products by comparing the complete chloroplast genome sequences of four *Zanthoxylum* species deposited in NCBI GenBank.

**Results:** Comparative analyses of the nucleotide diversity (*Pi*) of these *Zanthoxylum* genomes revealed four hypervariable divergent sites (*trnH-psbA, psbZ-trnG, trnfM-rps14*, and *trnF-ndhK*) with *Pi* > 0.025 among 520 windows. Of these four regions, including two genic and two intergenic regions, only *psbZ-trnG* yielded accurate PCR amplification results between commercial sancho and chopi products from the Korean herbal medicine market. We therefore selected *psbZ-trnG*, an InDel-variable locus with high discriminatory powers, as a candidate DNA barcode locus.

**Conclusion:** This InDel marker could be used as a valuable, simple, and efficient tool for identifying these medicinal herbs, thereby increasing the safety of these spices and herbal materials in the commercial market.

## Introduction

*Zanthoxylum* plants are thorny, deciduous shrubs and small trees with dense foliage, prickly trunks, and branches bearing edible fruits and leaves with a strong, pungent taste resembling lemon, anise, or mint. These plants, belonging to the Rutaceae family, comprise 250 species that are native to warm temperate and subtropical regions (Sun and Duan 1996; Epifano et al. 2011). While members of *Zanthoxylum* are commonly found in the Himalayan region, they also occur in Central, South, Southeast, and East Asia, and some species are found in America and Africa (Arun and Paridhavi 2012; Patiño et al. 2008; Appelhans et al. 2018). Many of these species are used as traditional medicines to treat human and animal diseases in Africa, Asia, and South America (Negi et al. 2011; Supabphol and Tangjitjareonkun 2014). Among these, *Z. schinifolium* (“sancho” in Korean, “huajiao” in Chinese) and *Z. piperitum* (“chopi” in Korean) are important traditional medicinal herbs in Korea and China (Kim et al. 2000; Ko and Han 1996; Yang 2008) that are also used as spices. *Korean Flora* describes seven *Zanthoxylum* species from Korea, including *Z. schinifolium* var. inermis (Nakai) T.B. Lee, *Z. piperitum* f. *pubescens* (Nakai) W.T. Lee, *Z. schinifolium* Siebold & Zucc., *Z. piperitum* (L.) DC., *Z. schinifolium* f. *microphyllum* (Nakai) W.T. Lee, *Z. planispinum* Sieblod & Zucc., and *Z. coreanum* Nakai (Lee 2003; Chung 1957; Nakai 1930). *Z. schinifolium* and *Z. piperitum* are the most ancient cultivated trees in southern regions of Korea. These trees are of major agricultural importance as sources of spices and traditional medicines (Ko and Han 1996; Yang 2008). Whereas *Z. schinifolium* thorns are arranged in an alternating pattern on branches, *Z. piperitum* thorns are arranged in an opposing pattern (Lee 2003). However, it is difficult to visually discriminate between dried and powdered herbal products made from *Z. schinifolium* and *Z. piperitum*. Furthermore, commercial herbal products commonly include a mixture of *Z. schinifolium* and *Z. piperitum* tissue. Since these plants are used in herbal medicines and as health food supplements, accurate methods should be developed to identify and characterize the two species.

Several studies have focused on developing molecular markers that can discriminate between various *Zanthoxylum* species used as traditional medicines in different countries. These molecular markers include amplified fragment length polymorphism (AFLP) markers for distinguishing between *Z. acanthopodium* and *Z. oxyphyllum* (Gupta and Mandi 2013), sequence-related amplified polymorphism (SRAP) markers (Feng et al. 2015), internal transcribed spacer (ITS) rDNA-specific markers (Kim et al. 2015), and ISSR markers (Feng et al. 2015). However, the use of these markers is limited by population dynamics and their reproducibility as diagnostic markers in the field. With the advancement of next-generation sequencing (NGS) technology, chloroplast (cp) genome assembly has become more accessible than Sanger sequencing. The development of molecular markers has also become cost effective through comparisons of cp genomes. The complete cp genomes of several *Zanthoxylum* species have been sequenced by de novo assembly using a small amount of whole genome sequencing data (Lee et al. 2016; Liu and Wei 2017; 2017).

In the current study, we compared the published cp genome sequences of four *Zanthoxylum* species and searched for species-specific regions with hypervariable nucleotide diversity among members of *Zanthoxylum*. Our aim was to develop interspecies insertion and deletion (InDel) markers that could be used to discriminate between *Z. schinifolium* and *Z. piperitum* and prevent the indiscriminate mixing of these two materials in commercial products.

## Results

### Comparative analysis of the cp genomes of various *Zanthoxylum* species

To investigate the level of sequence divergence among *Zanthoxylum* cp genomes, we performed a comparative analysis of three *Zanthoxylum* cp genomes with mVISTA using the annotated *Z. piperitum* sequence as a reference (Figure 1). The cp genomes of the *Zanthoxylum* species showed high sequence similarity, with identities of <90% in only a few regions, pointing to a high level of conservation among the cp genomes. However, sliding window analysis using DnaSP detected highly variable regions in the *Zanthoxylum* cp genomes (Figure 2). We calculated the nucleotide diversity (*Pi*) values in the four *Zanthoxylum* cp genomes as an indicator of divergence at the sequence level. The large single-copy (LSC) regions and small single-copy (SSC) regions were more divergent than the inverted repeat (IR) regions.

**Figure 1.**
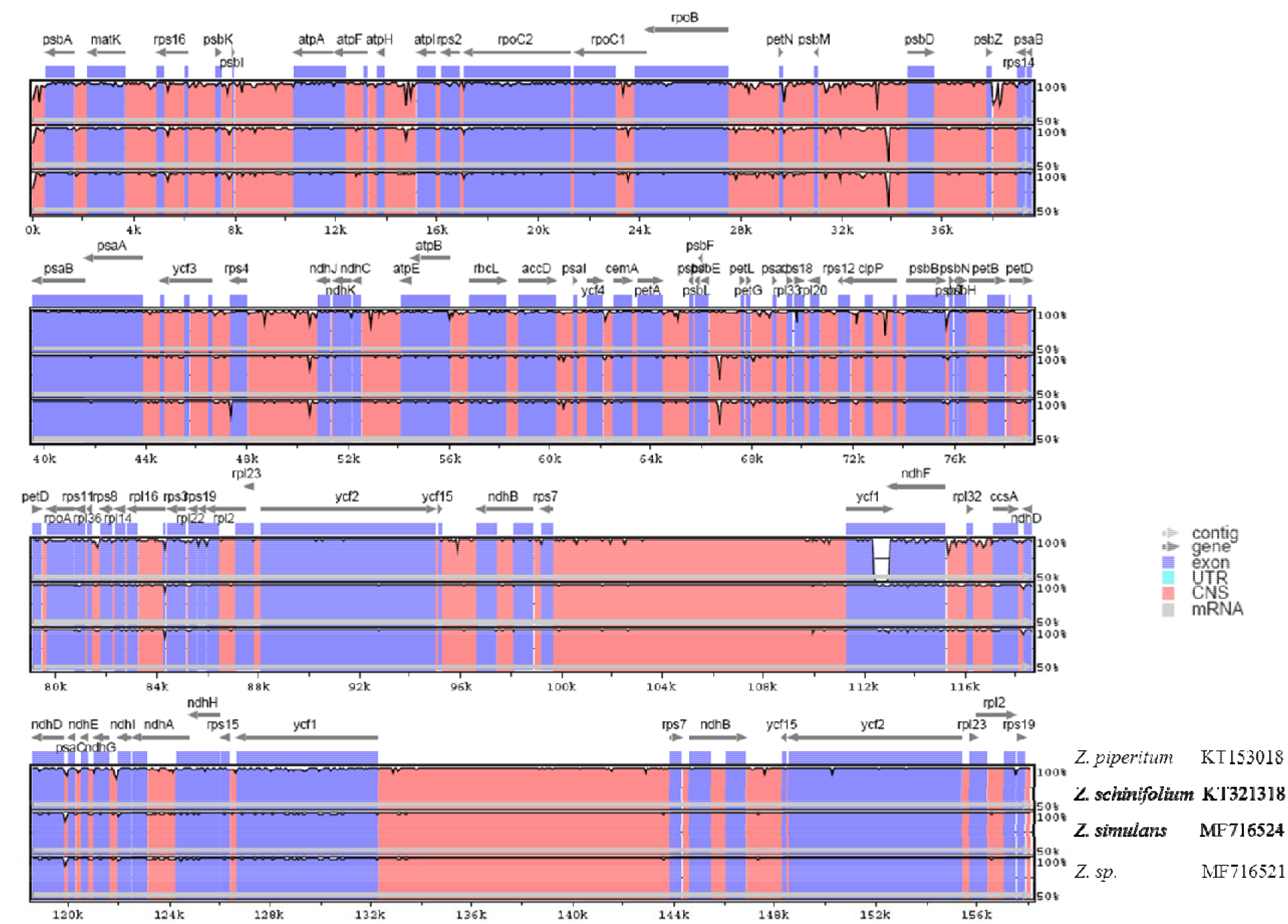
Comparison of the four *Zanthoxylum* cp genomes using mVISTA. The complete cp genomes of four *Zanthoxylum* species obtained from GenBank were compared, with *Z. piperitum* used as a reference. Purple blocks: conserved genes; pink blocks: conserved non-coding sequences (CNS). White represents regions with high levels of sequence variation among the four *Zanthoxylum* species.

**Figure 2.**
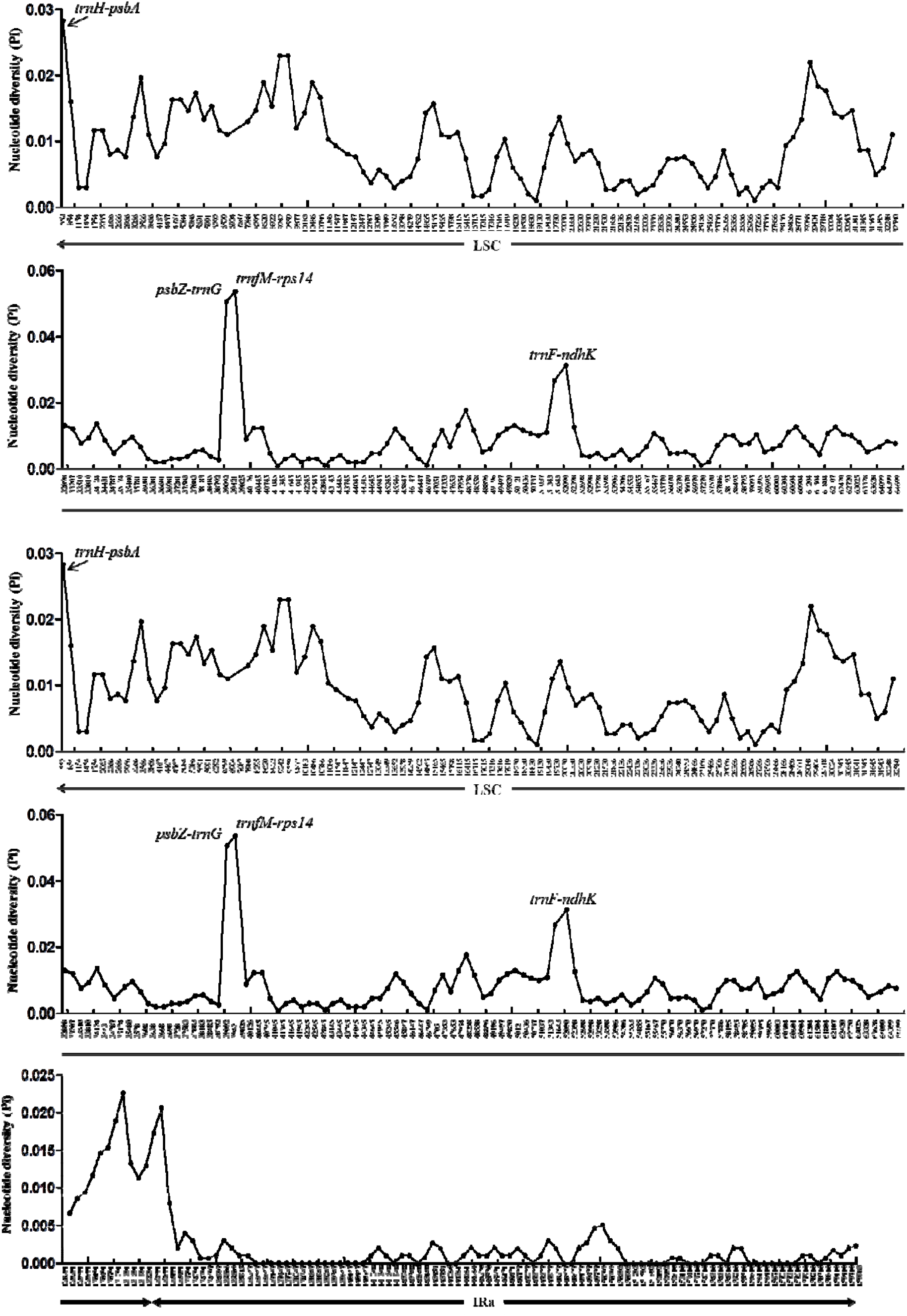
Sliding window analysis of the whole cp genomes of four *Zanthoxylum* species (window size: 600 bp; step size: 300 bp). X-axis: position of the midpoint of a window, y-axis: nucleotide diversity (*Pi*) of each window.

Among the 520 windows examined, IR regions were more conserved than LSC and SSC regions, with average *Pi* values of 0.0015 and 0.0012 for IRa and IRb, respectively (for regions other than those with a *Pi* value = 0). The *Pi* values for the LSC regions averaged 0.0082, whereas the SSC regions had a *Pi* value of 0.0103, and the average *Pi* value for all regions was 0.00609. Four mutational hotspots in the cp genomes showed markedly higher *Pi* values (>0.025), including two intergenic regions (*trnH-psbA*, 0.02833; *psbZ-trnG*, 0.05067) and two genic regions (*trnfM-rps14*, 0.05367; *trnF-ndhK*, 0.02667 and 0.03133) (Figure 2). Although the SSC and IR regions were generally highly conserved, the four regions located in the LSC regions were particularly divergent.

### Development of InDel markers to discriminate between *Z. schinifolium* and *Z. piperitum*

Based on multiple alignments of complete cp genome sequences, we selected the four most highly variable InDel loci as candidate DNA markers (Table 1). We confirmed these four InDel regions by PCR amplification and sequencing and investigated their utility for discriminating between *Z. schinifolium* and *Z. piperitum* (Figure 3). We produced four markers (ZanID1, ZanID2, ZanID3, and ZanID4) that were specific to *Z. schinifolium* and *Z. piperitum* and were derived from long InDels in the intergenic regions *psbZ-trnG, trnfM-rps14*, and *trnF-ndhK*, respectively. ZanID1 was derived from 11-, 3-, and 6-bp InDels in the *trnH-psbA* locus and was specific to *Z. schinifolium* and *Z. piperitum* (Figure 3). ZanID2 was derived from 19-, 28-, and 5-bp InDels and 40- and 23-bp tandem repeats (TRs) in the *psbZ-trnG* locus. ZanID3 was derived from 19-, 28-, 39-, and 6-bp InDels and a 23-bp TR in *trnfM-rps14*. ZanID4 was derived from 32-, 6-, 59-, and 50-bp InDels and an 8-bp TR in *trnF-ndhK*.

**Table 1.**
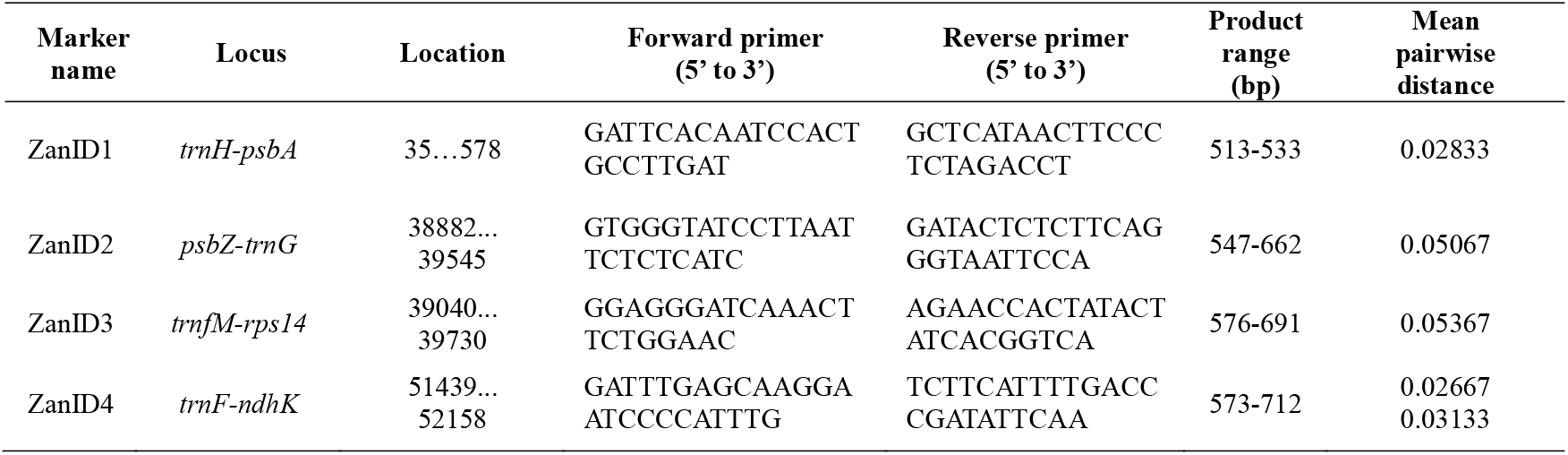
Primers used to amplify and sequence four highly variable loci

**Figure 3.**
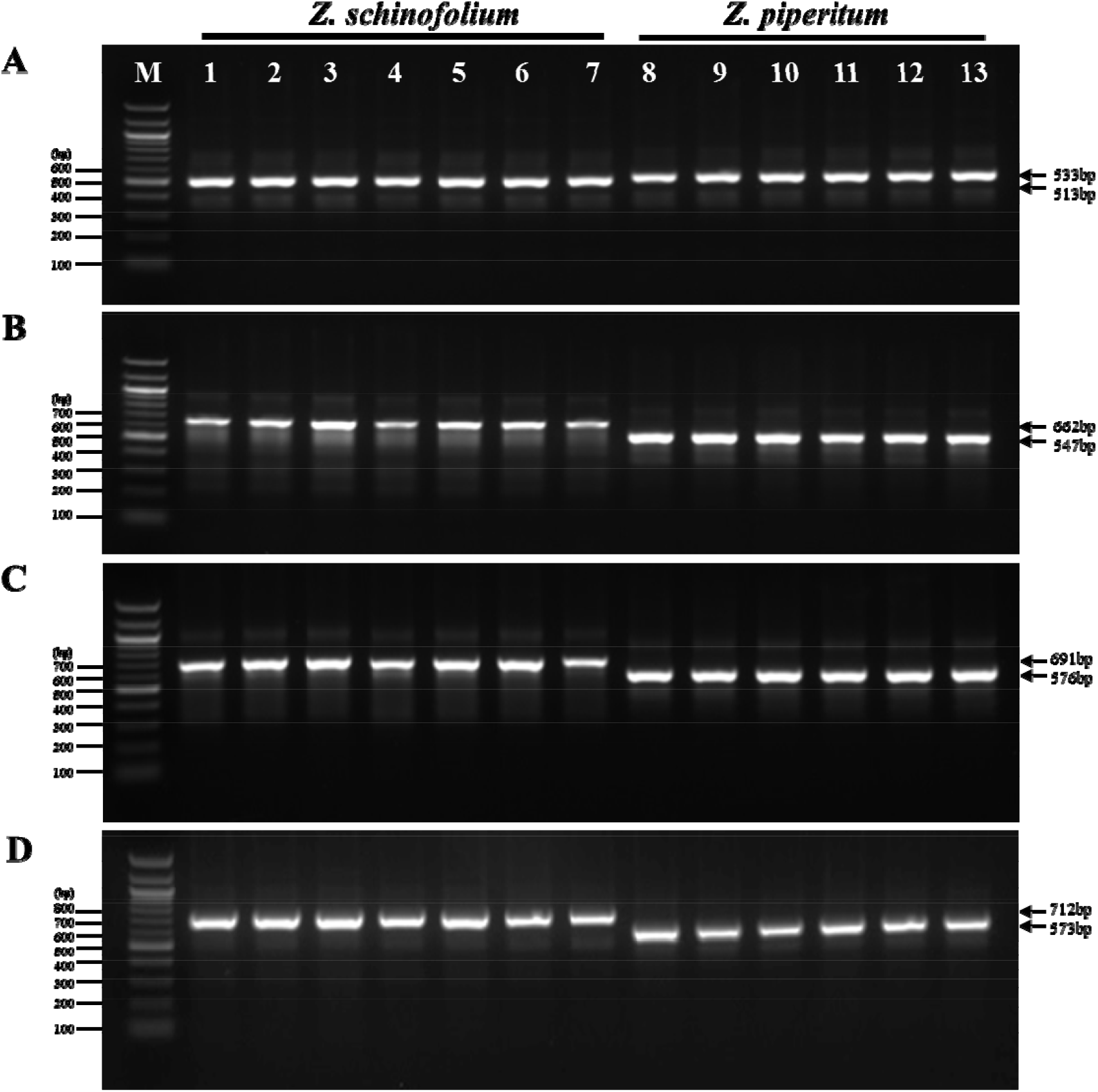
Gel profiles of fragments amplified from seven *Z. schinifolium* and six *Z. piperitum* samples using four pairs of primers derived from the *trnH-psbA* (A), *psbZ-trnG* (B), *trnfM-rps14* (C), and *trnF-ndhK* (D) loci in the four *Zanthoxylum* cp genomes. M: 100 bp ladder; 1– 7: amplicons from *Z. schinifolium* DNA; 8–13: amplicons from *Z. piperitum* DNA.

### Utilization of InDel markers in the Korean food market

To validate the utility of our newly developed markers to identify commercial dried herbal materials, we extracted genomic DNA from powdered or dried samples of seven sancho and six chopi products and amplified them by PCR using the newly developed primers (Table 1). The banding patterns of the *ZanID1* marker revealed that lanes 1, 2, 5, 6, 8, and 10 contained sancho samples, while lanes 3, 4, 9, and 11–16 were identified as chopi (Figure 4A). Different banding patterns were obtained using *ZanID2*, with lanes 2, 3, 5, 7, 8, and 10 containing sancho samples and the other samples identified as chopi (Figure 4B). Interestingly, samples 2, 5, and 7 clearly produced double bands in both species (Figure 4B). Furthermore, analysis of the banding patterns of the ZanID3 and ZanID4 markers revealed that lanes 2, 3, 5, 6, 7, 8, and 10 contained sancho samples, while the other samples were identified as chopi (Figure 4C and 4D). These three markers produced considerably different banding patterns, making it difficult to discriminate between sancho and chopi.

**Figure 4.**
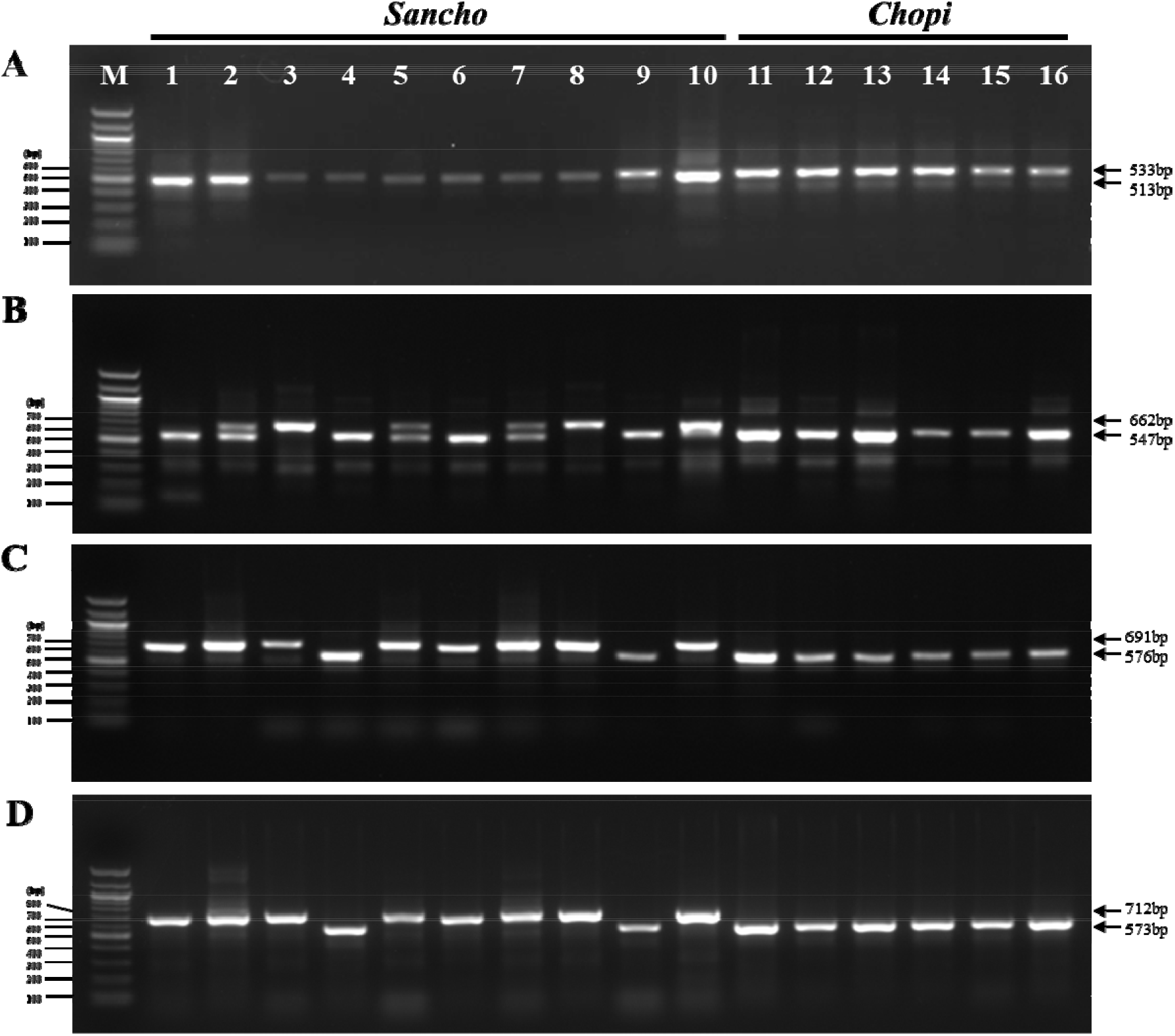
PCR identification of 16 commercial sancho and chopi products comprising dried or powdered seeds and pericarps. Lanes on 1.5% agarose gel; M: 100 bp ladder; 1–16: purchased commercial sancho and chopi products comprising dried or powdered seeds and pericarps (see Table 4 for details); 16 samples were amplified by PCR using four primer pairs; *ZanID1*_F and R (A), *ZanID2*_F and R (B), *ZanID3*_F and R (C), and *ZanID4*_F and R (D).

To select markers that could accurately discriminate between sancho and chopi, we performed PCR-RFLP analysis of the 13 samples by developing a PCR RFLP test to identify sancho samples. Many taxonomic studies of land plants have been based on the *trnH-psbA* region of cpDNA, as this DNA barcode exhibits high rates of sequence divergence among species. Based on the partial sequences of *trnH-psbA* in the cp genome that are shared between *Z. schinifolium* and *Z. piperitum*, we predicted that the PleI restriction enzyme would produce species-specific RFLP patterns and could therefore be used to identify *Z. schinifolium* based on the *trnH-psbA* locus. As shown in Figure 5, the fragment sizes for the two species were as follows: In *Z. schinifolium, PleI* produced two fragments (436 bp and 107 bp) from PCR products (lanes 2, 3, 5, 7, 8, and 10) of trnH-psbA (543 bp); in *Z. piperitum*, this enzyme did not digest the PCR products of trnH-psbA (562 bp). These results indicate that the psbZ-trnG marker is suitable for use as a reliable DNA barcoding tool to discriminate between *Z. schinifolium* and *Z. piperitum*.

**Figure 5.**
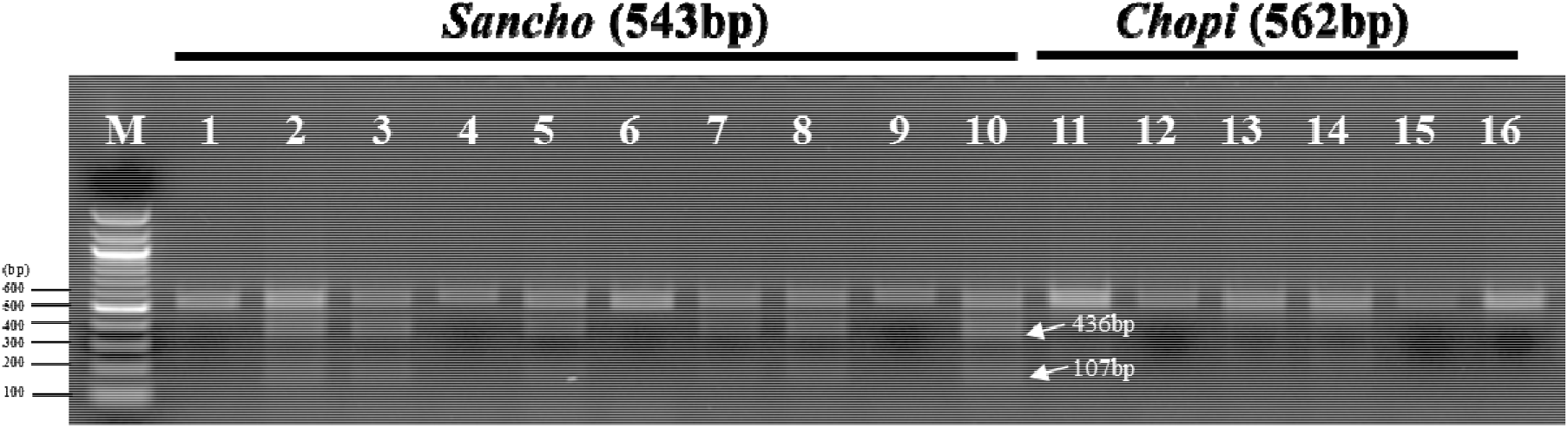
PCR-RFLP profiles of partial regions of *trnH-psbA* from all samples digested with *Ple*I. Numbers indicate sancho and chopi samples, as described in Table 4.

## Discussion

*Zanthoxylum* formed a phylogenetic group in previous molecular phylogenetic studies (Appelhans et al. 2018; Feng et al. 2015; Kim et al. 2015); however, these studies did not sufficiently resolve the relationships among some of its taxa. These studies were based on the ITS sequences of nuclear ribosomal DNA and the *trnL-trnF, matK-trnK, atpB, atp-rbcL*, and *rbcL* sequences of the cp genome (Sun et al. 2010; Feng et al. 2016; Zhao et al. 2018). Although these regions were considered to be universal DNA barcodes for higher plant, they did not allow to assess usefulness of these loci in barcoding of some taxa. Therefore, advances in NGS technologies have made it possible to sequence whole cp genomes and identify molecular markers. Highly variable markers derived from the cp genomes of different species at the genus level have uncovered many loci that are informative for systematic botany and DNA barcoding research (Dong et al. 2012; Li et al. 2015).

Here, we retrieved the complete cp genome sequences of four *Zanthoxylum* species from the NCBI database and compared species-specific cp diversity in *Zanthoxylum*. We confirmed the variation among species, with an average nucleotide diversity value (*Pi*) of 0.00609 among the four *Zanthoxylum* species. Although the average *Pi* value for the SSC region was relatively high (0.0103), high sequence divergence was detected at loci *trnH-psbA* (0.02833), *psbZ-trnG* (0.05067), *trnfM-rps14* (0.05367), and *trnF-ndhK* (0.02667 and 0.03133) in the LSC (0.0082). Indeed, the *trnH-psbA* locus is highly variable in most plants and is known as a universal DNA barcoding region (Kress et al. 2007; Whitlock et al. 2010). We used PCR amplification and sequencing to validate four hypervariable markers to distinguish between *Z. schinifolium* and *Z. piperitum* and to discriminate between sancho and chopi spice materials consumed in the online food market. However, PCR amplification of the four markers produced variable banding patterns, making it difficult to discriminate between sancho and chopi. Therefore, we designed a PCR-RFLP method using *PleI* digestion of the *trnH-psbA* DNA barcoding region, resulting in the production of two fragments (436 bp and 107 bp) only in *Z. schinifolium*. This marker, the ZanID2 marker from the *psbZ-trnG* region, is therefore suitable for discriminating between *Z. schinifolium* and *Z. piperitum* in sancho and chopi. The ZanID2 marker shows high sensitivity and specificity for detecting both sancho and chopi samples. Among the 10 sancho samples examined, three were successfully detected as sancho (30%), whereas three other samples (30%) produced a double band pattern that was clearly detected in both sancho and chopi samples. As expected, these results confirm the notion that products labeled as sancho that are sold in the spice and herbal medicine market often contain a mixture of sancho and chopi.

Although our results confirm that our newly developed InDel markers can be used to authenticate *Z. schinifolium* and *Z. piperitum* based on available cp genome data, more complete cp genome sequences are needed to comprehensively evaluate these InDel markers in the *Zanthoxylum* genus.

## Materials and methods

### Comparison of cp genomes and identification of hypervariable loci

All cp genome sequences in plants of the *Zanthoxylum* genus with complete genome sequence information were downloaded from GenBank (Table 1). The sequences were aligned using the Clustal W algorithm from MEGA 7.0 (Kumar et al. 2016) and CLC viewer 8.0 software. The gene distribution patterns and similarities in the *Zanthoxylum* cp genomes were compared and visualized using mVISTA software (http://genome.lbl.gov/vista/mvista/submit.shtml) in Shuffle-LAGAN mode with the annotated *Z. piperitum* KT153018 cp genome as a reference. The variability of the aligned genomes was evaluated using the sliding window method with DnaSP ver. 5.0. software (Librado and Rozas 2009). The window size was set to 600 base pairs (bp), the typical length of DNA markers. The step size was set to 300 bp for relatively accurate positioning of hypervariable InDels. Only regions with a nucleotide diversity (*Pi*) value of > 0.025 were considered. Hypervariable sites and genetic distance in the cp genome were calculated using MEGA 7.0. The InDel events were checked manually based on the aligned sequence matrix.

### Sample collection and genomic DNA isolation

Table 2 lists the *Z. schinifolium* and *Z. piperitum* collections from the National Institute of Biological Resources used in this study. Thirteen spice and powdered herbal materials (six sancho and six chopi samples) were purchased from verified local market sources in Korea and China (Table 4). Species identification was performed by the National Institute of Biological Resources, Korea. Prior to total genomic DNA extraction, 50 mg (dry weight) of each sample was added to a tube filled with stainless steel beads (2.38 mm in diameter) from a PowerPlant Pro DNA Isolation Kit (Qiagen, Valencia, CA), and the mixture was homogenized in a Precellys® Evolution homogenizer (Bertin Technologies). Genomic DNA was extracted from the collected samples using the PowerPlant Pro DNA Isolation Kit according to the manufacturer’s instructions.

**Table 2.** Complete cp genomes of the four *Zanthoxylum* species used in this study

| No. | Species | Research group | GenBank accession number | Sequence length (bp) |
| --- | --- | --- | --- | --- |
| 1 | *Z. schinifolium* | Crop Science and Biotechnology,<br>Department of Plant Sciences, Seoul<br>National University, Korea | KT321318 | 158,963 |
| 2 | *Z. piperitum* |  | KT153018 | 158,154 |
| 3 | *Z. simulans* | College of Forestry, Northwest A&F<br>University, China | MF716524 | 158,461 |
| 4 | *Z. sp.* |  | MF716521 | 158,572 |

**Table 3.**
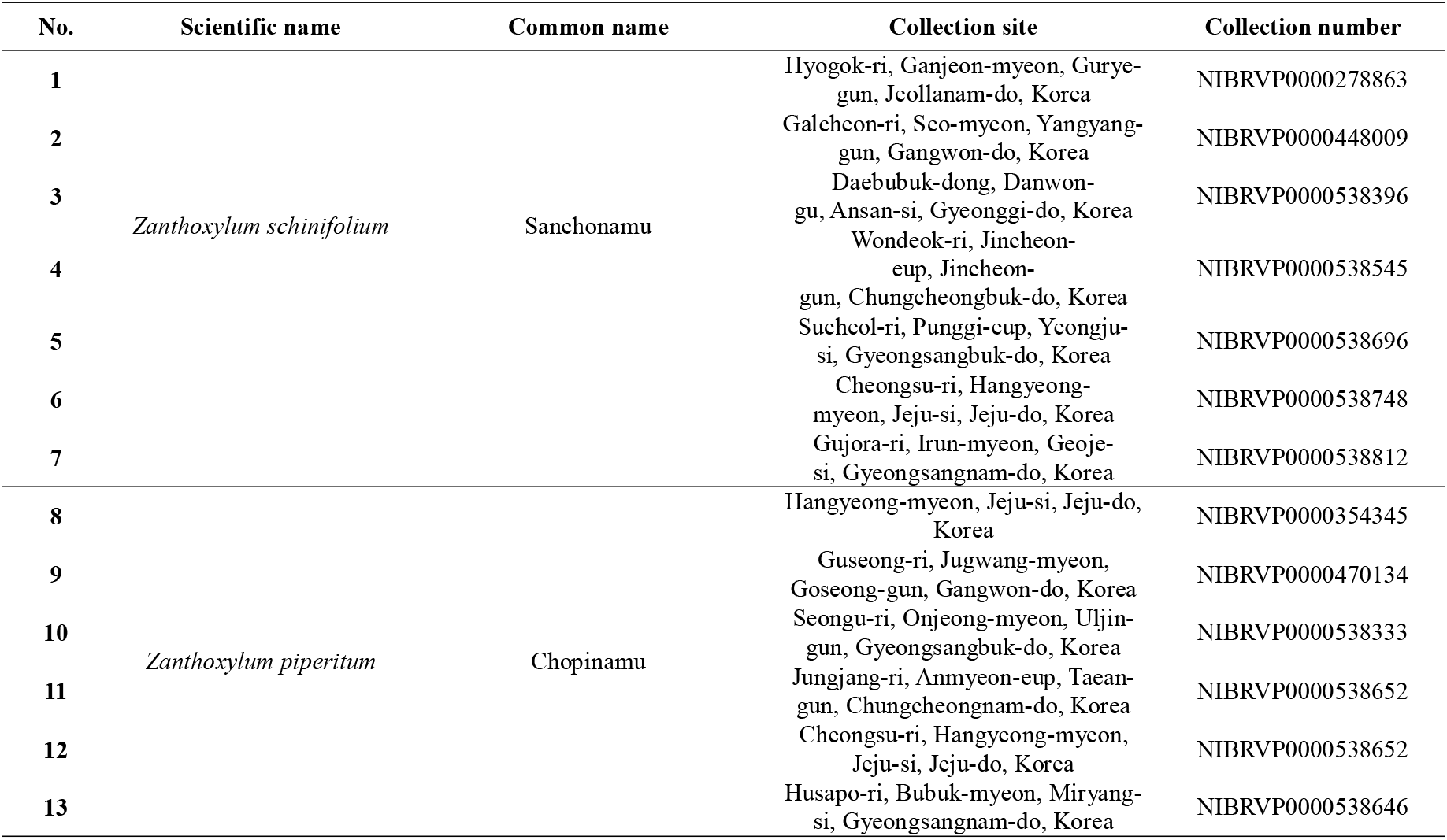
*Z. schinifolium* and *Z. piperitum* samples used in this study

| No. | Scientific name | Common name | Collection site | Collection number |
| --- | --- | --- | --- | --- |
| 1 | *Zanthoxylum schinifolium* | Sanchonamu | Hyogok-ri, Ganjeon-myeon, Gurye-gun, Jeollanam-do, Korea | NIBRVP0000278863 |
| 2 |  |  | Galcheon-ri, Seo-myeon, Yangyang-gun, Gangwon-do, Korea | NIBRVP0000448009 |
| 3 |  |  | Daebubuk-dong, Danwon-gu, Ansan-si, Gyeonggi-do, Korea | NIBRVP0000538396 |
| 4 |  |  | Wondeok-ri, Jincheon-eup, Jincheon-gun, Chungcheongbuk-do, Korea | NIBRVP0000538545 |
| 5 |  |  | Sucheol-ri, Punggi-eup, Yeongju-si, Gyeongsangbuk-do, Korea | NIBRVP0000538696 |
| 6 |  |  | Cheongsu-ri, Hangeong-myeon, Jeju-si, Jeju-do, Korea | NIBRVP0000538748 |
| 7 |  |  | Gujora-ri, Irun-myeon, Geoje-si, Gyeongsangnam-do, Korea | NIBRVP0000538812 |
| 8 | *Zanthoxylum piperitum* | Chopinamu | Hangeong-myeon, Jeju-si, Jeju-do, Korea | NIBRVP0000354345 |
| 9 |  |  | Guseong-ri, Jugwang-myeon, Goseong-gun, Gangwon-do, Korea | NIBRVP0000470134 |
| 10 |  |  | Seongu-ri, Onjeong-myeon, Uljin-gun, Gyeongsangbuk-do, Korea | NIBRVP0000538333 |
| 11 |  |  | Jungjang-ri, Anmyeon-eup, Taean-gun, Chungcheongnam-do, Korea | NIBRVP0000538652 |
| 12 |  |  | Cheongsu-ri, Hangeong-myeon, Jeju-si, Jeju-do, Korea | NIBRVP0000538652 |
| 13 |  |  | Husapo-ri, Bubuk-myeon, Miryang-si, Gyeongsangnam-do, Korea | NIBRVP0000538646 |

**Table 4.** The 16 commercial sancho and chopi spices and herbal products used in this study

| No. | Markets | Material component | Origin |
| --- | --- | --- | --- |
| 1 | A | Dried or powdered seeds and pericarps: sancho products (50–300 g) | China |
| 2 | B |  | China |
| 3 | C |  | China |
| 4 | D |  | Korea |
| 5 | E |  | China |
| 6 | F |  | China |
| 7 | G |  | China |
| 8 | H |  | China |
| 9 | I |  | Korea |
| 10 | J |  | Korea |
| 11 | K | Dried or powdered seeds and pericarps: chopi products (20–300 g) | Korea |
| 12 | L |  | Korea |
| 13 | M |  | Korea |
| 14 | N |  | Korea |
| 15 | O |  | Korea |
| 16 | P |  | Korea |

### Development and validation of the InDel molecular marker

To validate interspecies polymorphisms within the cp genomes and to develop DNA genetic markers for identifying these four *Zanthoxylum* species, primers were designed using Primer 3Plus (http://www.bioinformatics.nl/cgi-bin/primer3plus/primer3plus.cgi), and NCBI Primer-BLAST was performed based on the mutational hotspot regions (hypervariable regions) found in these *Zanthoxylum* cp genomes. PCR amplifications were performed in a reaction volume of 50 μL containing 5 μL 10x Ex Taq buffer (with MgCl_2_), 4 μL dNTP mixture (each 2.5 mM), Ex Taq (5 U/μL) (TaKaRa, Japan), 10 ng genomic DNA templates, and 1 μL (10 pM) forward and reverse primers. The mixtures were denatured at 95°C for 5 min and amplified for 40 cycles at 95°C for 30 s, 55°C for 20 s, and 72°C for 30 s, with a final extension at 72°C for 5 min. To detect PCR amplicons, the PCR products were separated by capillary electrophoresis (QIAxcel, Qiagen) using a QIAxcel DNA High Resolution Kit via the 0M500 method (Qiagen). The target DNA was extracted and purified using a MinElute PCR Purification Kit (Qiagen). Purified PCR products were sent to CosmoGenetech for sequencing (Seoul, Korea) with both forward and reverse primers. The sequencing results were analyzed by BLAST searches of the GenBank database. Sequence alignment and data visualization were carried out with CLC sequence viewer 8.0.

### RFLP analysis to identify suitable InDel markers

PCR was performed using universal primers for DNA barcodes within *trnH-psbA*: forward primer 5’-GTTATGCATGAACGTAATGCTC-3’ and reverse primer 5’-CGCGCATGGTGGATTCACAATCC-3’ (approximate product size: 401 bp). The PCR was conducted in a 50 μL reaction mixture containing 5 μL 10x Ex Taq buffer (with MgCl_2_), 4 μL dNTP mixture (each 2.5 mM), Ex Taq® (5 U/μL) (TaKaRa, Japan), 10 ng genomic DNA template, and 1 μL (10 pM) forward and reverse primers. The mixtures were denatured at 94°C for 5 min and amplified for 30 cycles of 94°C for 60 s, 55°C for 60 s, and 72°C for 90 s, with a final extension at 72°C for 7 min. The PCR-amplified products were separated by electrophoresis on 1.5% agarose gels for 30 min.

A 2-μL aliquot of each PCR-amplified *trnH-psbA* product (concentration 0.6 to 1 μg/ μL) was digested in 2 μL of 10x CutSmart buffer, 2 units of PleI restriction enzyme (New England Biolabs, Ipswich, MA; NEB), and 15.8 μL distilled H_2_O in a final volume of 20 μL, followed by incubation at 37°C for 20 min and inactivation at 65°C for 20 min. The digested fragments were separated by electrophoresis on 1.5% agarose gels stained with ethidium bromide, and the fragment patterns were visualized under UV light.

## Abbreviations

InDel: Insert and Deletion
RFLP: Restriction Fragment Length Polymorphism
CP: Chloroplast
LSC: Large single-copy
SSC: Smaill single-copy
IR: Inverted repeat
PI: pairwise distance

## Acknowledgments

This research was supported by the Support Program for Creative Industry Institutes (Commercial Biotechnology Sophistication Platform Construction Program, R0003950), funded by the Ministry of Trade, Industry & Energy (MOTIE, Korea).

## Authors’ Contributions

Y.K. and C.C. conceived and designed the experiments; S.-S.C. and Y.-P.H. collected the data; Y.K. and J.S. performed the experiments; Y.K. analyzed the data and wrote the manuscript. All authors read and approved the manuscript.

